# Recombinant Expression of Photo-crosslinkable 26S Proteasome Base Subcomplex

**DOI:** 10.1101/2024.12.16.628829

**Authors:** Santiago Yori Restrepo, Andreas Martin

## Abstract

The 26S proteasome complex is the hub for regulated protein degradation in the cell. It is composed of two biochemically distinct complexes: the 20S core particle with proteolytic active sites in an internal chamber and the 19S regulatory particle, consisting of a lid and base subcomplex. The base contains ubiquitin receptors and an AAA+ (ATPases associated with various cellular activities) motor that unfolds substrates prior to degradation. The *S. cerevisiae* base subcomplex can be expressed recombinantly in *E. coli* and reconstituted into functional 26S proteasomes *in vitro*, which allows the introduction of unnatural amino acids with novel functions or other mutations that may not be permissible *in vivo*. Here, we present a method for the introduction of the photo-induced crosslinking amino acid p-benzoyl-L-phenylalanine into the proteasomal base subcomplex. This approach has exciting implications for the study of protein-protein interactions of this complex that mediates the degradation of an incredibly diverse protein pool.

## Introduction

The 26S proteasome complex is the hub for regulated protein degradation in the eukaryotic cell. Its design as a compartmentalized protease, paired with the use of ubiquitin as a targeting signal and a complex network of ubiquitin ligases and deubiquitinating enzymes, ensures that cellular proteins are turned over selectively and in a tightly regulated manner [1]. The proteasome’s structural components reflect its task as a cellular machine to recognize and selectively degrade protein substrates. Degradation takes place in a compartmentalized peptidase, the 20S core particle (CP), which is composed of 14 distinct proteins, α1-7 and β1-7 [2]. The α subunits form a heptameric ring that rests atop a β heptameric ring. The complete 20S CP is assembled from four rings in an α_7_β_7_β_7_α_7_ layout, highlighting the role of the α rings as axial gatekeepers into the chamber that is formed by the β subunits. β1, β2, and β5 contain proteolytic active sites with trypsin, chymotrypsin, and caspase-like activities, respectively. Regulation of protein degradation is carried out by regulatory caps that bind to one or both sides of the 20S CP. Of these, the 19S regulatory particle (RP) is most prominent and best-characterized cap, forming the 26S proteasome holoenzyme [3]. The 19S RP can be biochemically divided into two subcomplexes, the base and the lid.

The base subcomplex is composed of 9 subunits. Six of these, Rpt1-6, form a heterohexameric motor of the AAA+ (ATPases Associated with cellular Activities) ATPase family [4]. This motor uses the energy of ATP hydrolysis to undergo conformational changes and generate the mechanical force required to unfold protein substrates and translocate the polypeptide into the 20S CP. Two of the non-ATPase subunits, Rpn1 and Rpn13, act as receptors for the ubiquitin degradation signal [5,6]. The lid and base subcomplexes of the *S. cerevisiae* 19S RP can be recombinantly expressed in *E. coli* and *in vitro* reconstituted with endogenous 20S CP, allowing the generation of mutant 26S holoenzyme that otherwise would not support cell viability [4]. This also extends to the ability of leveraging methodologies of genetic code expansion to introduce unnatural amino acids (UAAs) with novel functions, as was previously shown with the incorporation of the UAA p-azido-L-phenylalanine (AzF) in amber stop-codon suppression [7]. The addition of AzF to either the lid or base subcomplex required the development of a plasmid encoding an AzF tRNA synthetase/tRNA pair to be coexpressed in *E. coli* together with an existing system for the expression of proteasome subunits and assembly factors. This system was originally used for attaching fluorescent probes to decipher the kinetic pathways of the proteasome in bulk assays [8], and is now additionally employed for single-molecule fluorescence and FRET-based experiments [9-11].

Here, we describe the incorporation of a different UAA for the purpose of site-specifically crosslinking protein substrate to the base of the 19S RP. Similar to the incorporation of AzF, tRNA/tRNA synthetase pairs for the incorporation of the photo-induced crosslinking UAA p-benzoyl-L-phenylalanine (Bpa) have been developed [12,13]. Cross-linking is often used for structural and functional studies, and Bpa incorporation therefore has great potential for furthering our understanding of the 26S proteasome. Bpa was developed for the purpose of crosslinking and has exceptional chemical features. Its aryl ketone group is chemically stable, can be manipulated in ambient light, is excited into a diradical triplet state with low-energy UV light (365 nm), does not readily react with water in solution, and the probe can be repeatedly reactivated if no initial cross-linking occurs (**Figure 1**). In contrast to traditionally used crosslinkers in solution that react non-specifically with nucleophilic amino acids, the incorporation of Bpa results in crosslinks to a specific site on the protein—only where the suppressed amber codon was introduced. This is especially useful for large multi-protein complexes with many solvent-exposed, potentially crosslinkable lysine and cysteine residues, such as the proteasome. Additionally, the light-induced activation of Bpa’s benzophenone allows for exceptional temporal control of crosslinking. Its incorporation required the generation of a new plasmid for recombinant expression.

**Figure 1:**
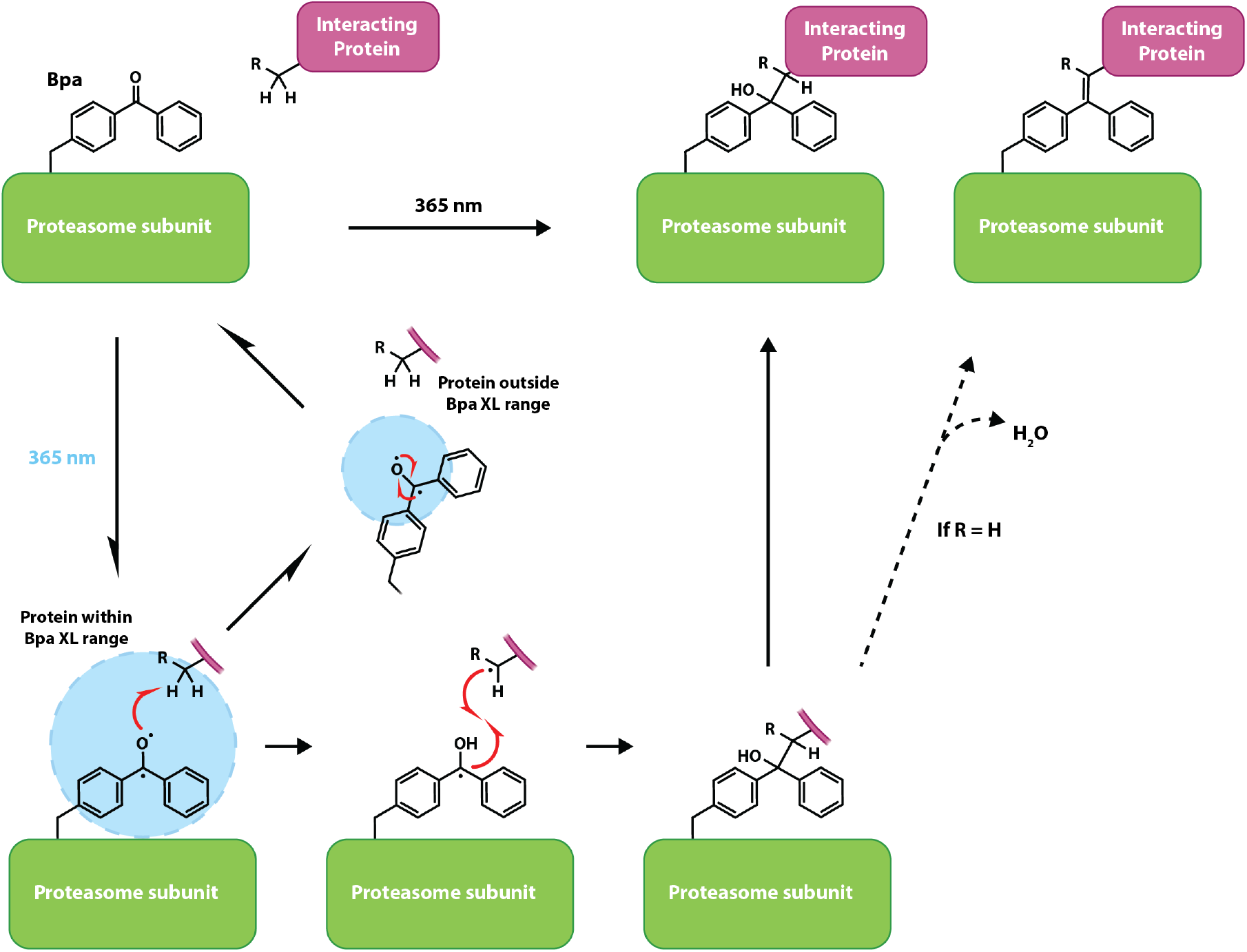
Photo-induced crosslinking of Bpa. Bpa (p-benzoyl-L-phenylalanine) is a crosslinking unnatural amino acid. Upon irradiation by 365 nm light, the ketone of the benzophenone sidechain is excited to a triplet state, which allows it to react with carbon-hydrogen bonds within its crosslinking range. If there is no suitable crosslinking position within range, the triplet state collapses and reforms the Bpa, which can undergo subsequent rounds of excitation and crosslinking. C-H bonds within the radius will undergo hydrogen abstraction and then bond formation between the carbonyl carbon of the Bpa and the target to form a benzopinacol-like covalent adduct. If the sidechain is a glycine, the newly formed covalent adduct can undergo rearrangement and loss of water to form an alkene [14].

To this end, the tRNA-synthetase pair from the pEVOL-Bpa plasmid developed by the Schultz lab [13] was subcloned into a pULTRA backbone. This new plasmid, pULTRA-Bpa, resulted in IPTG-inducible expression of the tRNA/tRNA-synthetase pair to complement the existing plasmids for the recombinant expression proteasomal subunits (**Figure 2A**).

**Figure 2:**
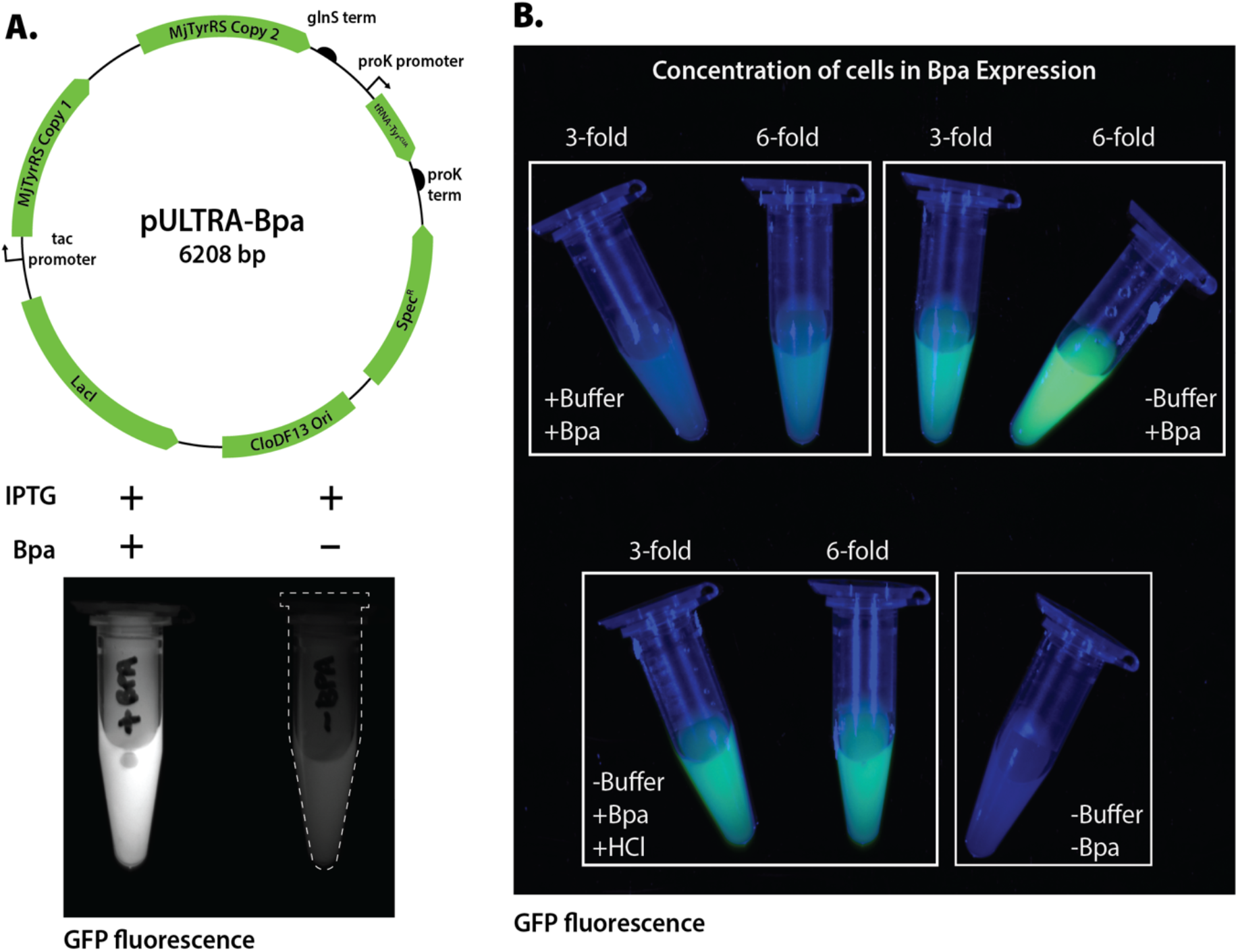
A pULTRA-Bpa plasmid for the expression of Bpa-containing proteins. Bpa can be site-specifically introduced into a protein using amber stop codon suppression. **A**. Top, plasmid map for the pULTRA-Bpa plasmid used in this protocol shows the tac promoter expression of two copies of the tRNA synthetase for Bpa incorporation. Bottom, expression test with an amber codon containing GFP test construct for the IPTG-inducible expression with the Bpa incorporation machinery. Amber suppression is shown to be Bpa-dependent. **B**. Bpa incorporation is dependent on its availability to the cell. Expression tests using the same GFP reporter construct show that concentrating cells 6-fold allows better exposure to the Bpa for incorporation. Conversely, addition of a buffering agent to the Terrific Broth media resulted in precipitation of the Bpa and poor incorporation. Bpa is dissolved in 1 M NaOH, and neutralization of the media with 1 M HCl after addition of the Bpa results in reduced incorporation. Culture ODs were normalized to the -Buffer -Bpa control. GFP fluorescence images of cell cultures were captured on a ChemiDoc MP using the fluorescein 532/30 filter set and automatic exposure.

The most expensive and limiting reagent in a UAA-containing protein expression experiment is the UAA itself, which is typically added to a final concentration of 2 mM. In order to maximize delivery of the UAA into cells, a large culture is grown in a complete media until reaching approximately mid-log phase, before cells are harvested by centrifugation and resuspended in a smaller volume of buffered Terrific Broth (TB) supplemented with the UAA. The higher cell density after this concentration step maximizes the use of the UAA reagent. Previous work with AzF incorporation used a six-fold concentration of cells into buffered TB media [7]. The solubility of Bpa required dissolving it in sodium hydroxide, and systematic testing was carried out to determine the ideal conditions for base expression and Bpa incorporation. In order to rapidly optimize growth conditions, we used a GFP reporter with an amber TAG codon positioned such that the fluorescence only evolves upon proper incorporation of Bpa. We determined that a 6-fold concentration of cells results in robust Bpa incorporation. However, unlike with AzF incorporation, the addition of a buffering component into the TB media resulted in precipitation of the Bpa, making it unavailable to the *E. coli* cells and resulting in poor GFP incorporation, as assayed by fluorescence (**Figure 2B**). Other groups have reported neutralizing the NaOH after dissolving the Bpa through the addition of an equivalent amount of HCl, which we found to also reduce Bpa incorporation due to precipitation. After concentration, the cells are briefly incubated in TB with Bpa to allow absorption of the UAA. Expression of the recombinant proteasome subcomplex and the UAA machinery is then induced by the addition of IPTG. As a proof-of-concept, we recombinantly expressed the base subcomplex with Bpa incorporated in the ATPase subunit Rpt1.

The purification of the base described below is carried out by a two-step affinity procedure: binding of the 6x-histidine tag on Ni-NTA resin and elution with imidazole, followed by binding of a secondary FLAG tag to anti-FLAG antibody resin and elution with FLAG peptide. As a last step, the complex is buffer exchanged and further purified by size-exclusion chromatography (SEC). To verify that the 280 nm UV detector of the FPLC instrument did not affect the stability of the Bpa during purification, the SEC step was carried out with and without the UV detector, which revealed that exposure to the detector light did not induce unwanted crosslinking. We further tested whether incorporation of Bpa altered the activity of the proteasome by testing the ATPase rates of the base subcomplex. The Bpa-containing base subcomplex exhibited an ATPase activity like that of the unmodified subcomplex, with a rate of ∼ 40 ATP min^-1^ (**Figure 3A**). Finally, we tested whether the Bpa in Rpt1 would internally crosslink to other subunits within the base subcomplex or even Rpt1 itself upon exposure to 365 nm UV light for 15 minutes. We observed that a crosslinked product appeared only after UV exposure (**Figure 4B**), which we identified as the adjacent motor subunit, Rpt2.

**Figure 3:**
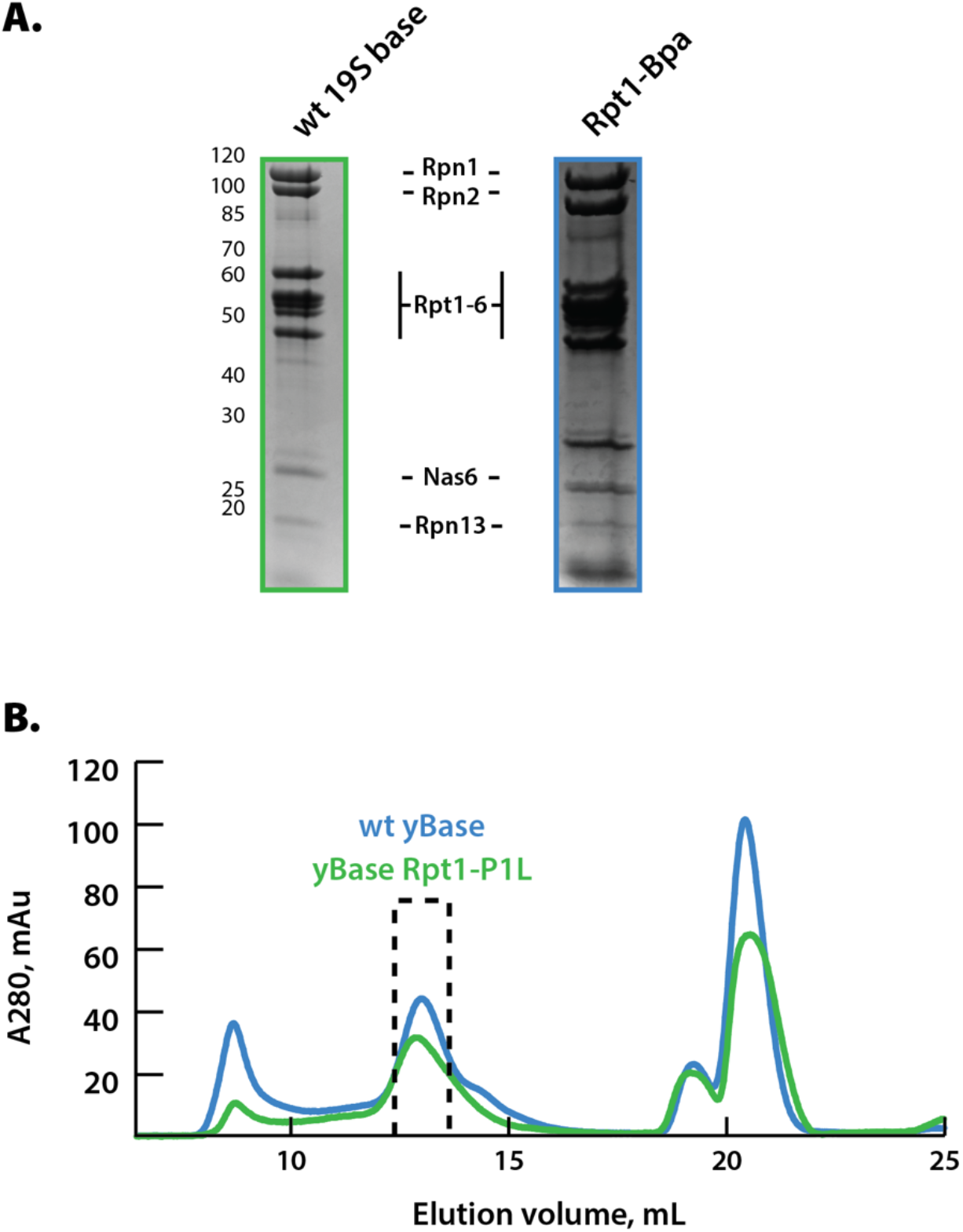
Purification of Bpa-containing 19S RP base subcomplex. **A**. Coomassie-stained SDS-PAGE comparison of unmodified and Bpa-containing base complexes. **B**. Size-exclusion chromatography trace for a Superose 6 Increase 10/300 column. Dotted box represents fraction that were concentrated for PAGE gels shown in A.

**Figure 4:**
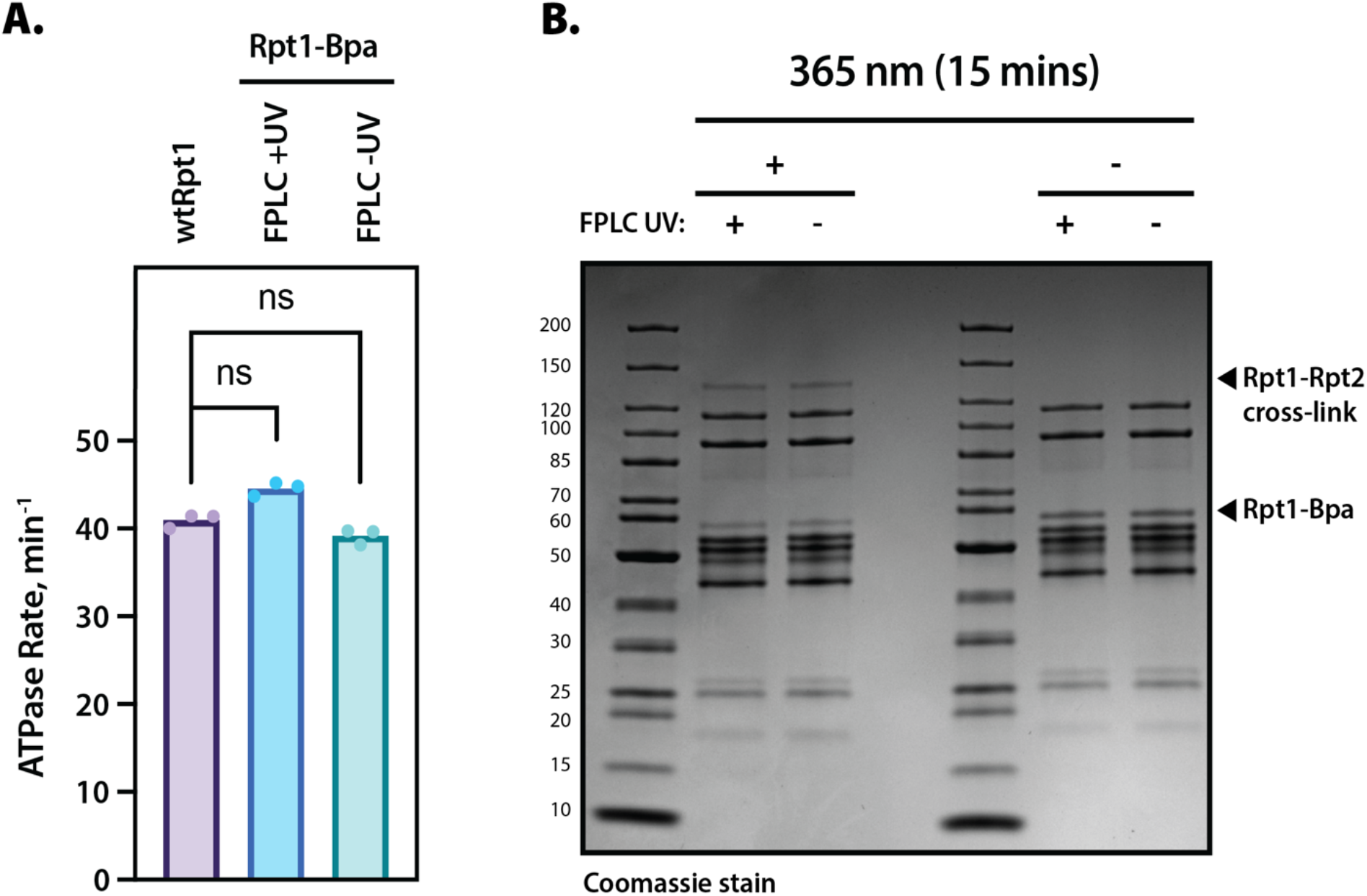
ATPase and Crosslinking Assays Demonstrate Dependence of Bpa-Base crosslinking on UV exposure. **A**. ATPase rate of recombinant wild-type and Bpa-containing 19S RP base complex are comparable. Bpa-containing base complex was purified as described, whereby the SEC step was performed with the 280 nm UV detector activated or deactivated to test for detector-induced unwanted crosslinking. Individual measurements and their mean are shown for N= 3 technical replicates. Statistical significance was calculated using a ratio paired t-test. **B**. Crosslinking assay visualized by Coomassie-stained SDS-PAGE gel demonstrates that the Bpa-containing base complex crosslinks only upon irradiation by 365 nm light and not from the 280 nm detector of the FPLC.

As the site of mechanical substrate processing, ubiquitin recognition, and cofactor binding, the base subcomplex of the 19S RP provides an exciting platform for the introduction of a site-specific photo-induced crosslinker, and our protocol thus describes the important expansion of the toolkit for probing proteasome function.

## Materials

### Media

1. Lysogeny Broth (LB) agar (1L): 10 g tryptone, 5 g yeast extract, 10 g NaCl, 15 grams agar.
2. dYT Media (1 L): 16 g tryptone, 10 g yeast extract, 5 g NaCl, add 20 g agar for plates.
3. Unbuffered Terrific broth (TB) Media (1 L): 12 g tryptone, 24 g yeast extract, 4 mL glycerol.

### Reagents

4. 1000X ampicillin: 300 mg/mL ampicillin sodium (GoldBio) in 50% ethanol.
5. 1000X kanamycin: 50 mg/mL kanamycin monosulfate (GoldBio) in water.
6. 1000X chloramphenicol: 25 mg/mL chloramphenicol (Thermo-Fisher) in 100% ethanol.
7. 1000X spectinomycin: 100 mg/mL spectinomycin dihydrochloride pentahydrate (GoldBio) in water.
8. 1 M Isopropyl β-D-1-thiogalactopyranoside (IPTG) (GoldBio).
9. 1 M NaOH: 20 g in 500 mL, filter sterilize.
10. 4-benzoyl-L-phenylalanine (Amatek).
11. 1000X Aprotinin (GoldBio): 1 mg/mL in 25% methanol.
12. 1000X Pepstatin (GoldBio): 1 mg/mL in 90% methanol, 10% acetic acid.
13. 1000X Leupeptin (GoldBio): 1 mg/mL in water.
14. 100X AEBSF (GoldBio): 100 mM in water.
15. Benzonase nuclease (Novagen).
16. Lysozyme (GoldBio).
17. 2 M imidazole: 68.08 g in water, adjust pH to 7.6, top to 500 mL, filter sterilize.
18. 1 M HEPES: 119.15 g in water, adjust pH to 7.6, top to 500 mL, filter sterilize.
19. 1 M Tris HCl: 60.57 g in water, adjust pH to 7.5, top to 500 mL, filter sterilize.
20. 2 M MgCl_2_: 101.65 in water, top to 250 mL, autoclave.
21. 0.5 M ATP (VWR): Dissolve 23 grams in 60 mL water. Adjust pH dropwise with NaOH until 7.0-7.4, bring the volume to 75 mL, make 1 mL aliquots and store at - 80°C.
22. 0.5 M TCEP (GoldBio): Add 5.73 g tris(2-carboxyethyl)phosphine-HCl to 30 mL of water. Adjust pH to 7.0 with 5 M NaOH and adjust volume to 40 mL with water. Aliquot and store at −20 °C.
23. 3X FLAG (MDYKDHDGDYKDHDIDYKDDDDK) peptide (GenScript): Dissolve peptide in Tris-Buffered Saline (TBS), adjust pH to 7.6 using NaOH.
24. Bradford Reagent.
25. 20X ATPase Assay Mix: 375 U/mL pyruvate kinase (Sigma), 429 U/mL lactate dehydrogenase (Sigma), 20 mM NADH (Sigma), 150 mM phosphoenolpyruvate (Sigma), 50 mM ATP (VWR) in GF Buffer at pH 7.6.

### Buffers

26. NiA Buffer: 60 mM HEPES, 100 mM NaCl, 100 mM KCl, 10 mM MgCl_2_, 20 mM imidazole, 5% glycerol, adjust pH to 7.6.
27. NiB Buffer: NiA with 250 mM imidazole, re-adjust pH to 7.6.
28. FLAG Regeneration Buffer: 0.1 M glycine, adjust pH to 3.5 with concentrated HCl.
29. Tris-Buffered Saline (TBS): 50 mM Tris HCl, 150 mM NaCl, pH 7.5.
30. Gel-filtration (GF) buffer: 30 mM HEPES, 50 mM NaCl, 50 mM KCl, 10 mM MgCl_2_, 5% glycerol, adjust pH to 7.6, filter sterilize and de-gas prior to use.

### Resins

31. Ni-NTA agarose (Sigma).
32. Anti-FLAG M2 agarose (Sigma).

### Other materials

33. Bio-Rad Micropulser Electroporator.
34. Bio-Rad 0.2 cm electroporation cuvettes.
35. pULTRA-Bpa plasmid.
36. Probe-tip sonicator.
37. Stainless steel beaker.
38. Corning 384-well low-volume clear bottom plate.
39. Superose 6 Increase 10/300 SEC chromatography column (Cytiva).
40. Blak-Ray B-100AP 365 nm lamp (Thermo-Fisher).
41. Bio-Rad Mini-PROTEAN TGX 4-20% polyacrylamide gel.
42. 2X sample buffer (Bio-Rad).

## Methods

### Expression of Bpa-containing Base subcomplex

1. Electroporate the pULTRA-Bpa plasmid into electrocompetent BL21(DE3) containing the three 19S RP base expression plasmids, as previously described [7] (*see* **Note 1**). Plate on an LB plate containing 1X each of ampicillin, kanamycin, chloramphenicol, and spectinomycin.
2. Grow overnight at 37 °C.
3. From the plate, pick a single colony and start a 5 mL overnight culture in dYT with 1X of all four antibiotics.
4. Make two overnight cultures by adding 1 mL of the 5 mL cultures to 50 mL dYT with 1X antibiotics and grow overnight at 37 °C.
5. The following day, transfer the 50 mL cultures into 50 mL conical tubes and spin cells down at 4,000xg for 10 minutes.
6. Wash the cell pellets by resuspending each in 30 mL of fresh dYT.
7. Centrifuge and repeat the wash step once more.
8. Inoculate 6L of dYT with 0.5X antibiotics in 1L baffled flasks by adding 10 mL of the cell suspension to each flask (*see* **Note 2**).
9. Grow at 37 °C and monitor the OD_600_.
10. While the cells are growing or prior to growth, autoclave and cool down 1 L centrifuge bottles.
11. Once the cells reach an OD_600_ of 0.6, harvest the cells by centrifuging at 4000xg for 15 minutes.
12. Resuspend the cells in 1 L of unbuffered TB and split into two flasks to maximize oxygenation of the media.
13. Dissolve 538.6 mg Bpa in 300 μL 1 M NaOH and divide between the two flasks to reach a final concentration of 2 mM Bpa in each flask (*see* **Note 3**).
14. Shake cells for 30 minutes to allow uptake of the Bpa and monitor the OD_600_ to make sure the cells are growing.
15. After this incubation, induce expression of the 19S RP base and the UAA tRNA/synthetase pair by adding 0.5 mL of 1 M IPTG (1 mM final) to each flask.
16. Express protein for 5 hours at 30 °C, followed by 16 °C overnight.
17. Harvest the cells by centrifuging at 4,000xg for 15 minutes.
18. Resuspend the pellets with cold NiA buffer with added 1X protease inhibitors and 50 U of benzonase such that the final volume is approximately 75 mL.
19. Transfer the cell suspension to two 50 mL conical tubes, flash-freeze in liquid nitrogen, and store at -80 °C or proceed with the purification protocol below.

### Purification of Crosslinkable 19S RP Base Subcomplex

1. Thaw the tubes in water and transfer the contents to a 250 mL stainless steel beaker. Add 100 μg/mL of lysozyme.
2. Sonicate the lysate on ice using a probe-tip sonicator set to 70% amplitude, with a cycle of 3 seconds on and 59 seconds off.
3. Clarify the lysate by centrifugation at 26,000xg for 30 minutes at 4 °C. **All subsequent steps should be carried out in a 4 °C cold room**.
4. While the lysate is centrifuging, measure 10 mL of 50% Ni-NTA agarose slurry and add to a 50 mL conical tube. Spin the slurry at 700xg for 3 minutes to settle the agarose.
5. Resuspend the resin by topping off the conical tube with water to wash off the storage solution, spin to settle.
6. Repeat the wash step once more.
7. Carefully decant the water and resuspend the agarose with NiA buffer to 20 mL.
8. Spin to settle and repeat the resuspension.
9. Transfer the clarified lysate to two clean 50 mL conical tubes. Add 10 mL of Ni-NTA agarose in NiA buffer to each conical tube.
10. Batch-bind the lysate to the Ni-NTA at 4 °C with rotation for up to one hour.
11. Prepare a glass gravity chromatography column by washing with water and then NiA buffer. Transfer the batch-bound mixture to the column and allow the resin to settle.
12. Open the valve and allow the lysate to flow through. **All subsequent steps require buffers supplemented with 1 mM ATP**.
13. Wash the resin with 25 mL of NiA buffer + 1 mM ATP.
14. Repeat the wash step.
15. Elute the base complex by adding 20 mL of NiB buffer + 1 mM ATP. Collect the elution in a 50 mL conical tube.
16. Regenerate two 5 mL FLAG columns:
  a. Add 10 mL of TBS and gently resuspend the resin, open the column valve and allow the buffer to flow and the resin to re-pack.
  b. Wash once more with 10 mL of TBS, close the valve when the buffer is just above the resin. **Do not let the resin run dry**.
  c. Regenerate the resin by flowing three sequential 5 mL volumes of 0.1 M glycine regeneration buffer. It is important to monitor the resin at this step, so it does not run dry or sit in the glycine buffer for more than 20 minutes.
  d. Wash the resin with 10 mL TBS.
  e. Equilibrate the resin with 10 mL of NiA buffer + 1 mM ATP. The resin is now ready for binding the protein from the Ni-NTA elution.
17. Split the Ni-NTA elution sample across the two FLAG columns. Allow the sample to flow through the FLAG resin and collect the flowthrough in 15 mL conical tubes.
18. When the volume approaches the resin bed, close the valve and top off the resin with the flowthrough. Repeat this binding step 4 times, ensuring the sample has been in contact with the resin for at least 30 minutes.
19. Wash each FLAG column with 25 mL of NiA buffer + 1 mM ATP.
20. Repeat the wash.
21. While the columns are washing, prepare two conical tubes with 15 mL of NiA buffer + 1 mM ATP and 0.15 mg/mL 3X FLAG peptide.
22. Add this elution buffer to the column, allow it to flow through, and collect the elution.
23. Pre-equilibrate a 15 mL 100K MWCO concentrator by adding NiA buffer + 1 mM ATP and spinning at 4°C.
24. Add the FLAG elution sample and concentrate at 4 °C to approximately 500 μL by spinning at 2,000xg in 5 minute increments.
25. Filter the concentrated protein with a pre-equilibrated 0.22 μm spin filter.
26. Equilibrate a Superose 6 Increase 10/300 size-exclusion chromatography column with GF buffer + 0.5 mM ATP and 0.5 mM TCEP.
27. Inject the protein concentrate into the FPLC using a 0.5 mL loop and run, making sure to collect 0.5 mL fractions. The base subcomplex should elute at an elution volume of approximately 13 mL (*see* **Figure 3B**). Collect the fractions corresponding to the eluted base subcomplex.
28. Concentrate using a 500 μL 100K MWCO concentrator to approximately 100 μL.
29. Take a small sample for SDS-PAGE analysis and for measuring protein concentration by Bradford assay.
30. Aliquot and flash freeze samples in liquid nitrogen. They can be stored at -80 °C, but will not tolerate subsequent freeze-thaw cycles.

### ATPase Assay

1. Pre-warm a plate reader and a 384-well plate to 30 °C at least 15 minutes before making a measurement.
2. Calculate the volume needed and dilute purified base complex to prepare a 2X enzyme mixture in a 0.65 mL Eppendorf tube. For example, if measuring the ATPase rate of 150 nM of base, a 2X enzyme mix will need 300 nM of base. Aliquot 7 μL of the 2X enzyme mix into a labelled PCR strip in triplicate and incubate on ice.
3. Dilute 20X ATPase mixture to 2X in another Eppendorf tube on ice. Aliquot 10 μL of this ATPase mix into a separate PCR strip.
4. Incubate both mixes at 30°C for 3 minutes in a thermocycler or a heat block.
5. Using a multichannel pipette, transfer 7 μL of the ATPase mix into the enzyme PCR strip. Mix by carefully pipetting up and down.
6. Using a separate multichannel pipette, transfer 10 μL of the 1X enzyme/ATPase mixture into the wells of the pre-heated 384-well plate.
7. Monitor the NADH absorbance at 340 nm with pathlength correction for 30 minutes.
8. Calculate the ATP hydrolysis rate from the slope of the loss of absorbance at 340 nm.

### Crosslinking of Bpa-containing Base complex

1. Dilute purified base complex into a PCR strip to the desired concentration and a final volume of 50 uL (*see* **Note 4**).
2. Incubate the PCR strip at 4 °C either with a chilled PCR aluminum block or with packed ice.
3. Remove the PCR strip caps and place a 365 nm UV lamp 2 centimeters from the openings of the strip (*see* **Note 5**).
4. Expose the crosslinkable base complex to the light for 15 minutes, maintaining at 4 °C.
5. Add 2X sample buffer to the samples, run on an 4-20% Mini-PROTEAN TGX SDS polyacrylamide gel, and visualize using Coomassie stain to assess crosslinking (*see* **Note 6**).

### Notes

1. It is important to prepare fresh plates with quadruple antibiotics and make fresh transformations of cells with all four plasmids. The four antibiotics have different shelf lives and the cells readily lose selection plasmids. It is quite common to pick a colony from an old plate and find that it does not grow on fresh media with four antibiotics. An option is to carry out a colony PCR from the plate or starter culture to ensure cells contain the four plasmids prior to moving forward with expression.
2. It is possible to grow the cells in 1X antibiotics for this step, but we have found that the cell growth is quite slow in these conditions. A concentration of 0.5X of antibiotics results in a growth rate that is more amenable to a day’s experiment while still reducing cell contamination.
3. The 300 μL volume of 1 M NaOH was empirically determined to be a minimal amount needed to dissolve the Bpa while still preventing precipitation upon addition to the media. The quality of Bpa is batch dependent and it is recommended to sterilize the solution by passing through a 0.22 μm filter prior to adding to the cultures, as the powdered Bpa can sometimes be contaminated with bacteria.
4. Initial crosslinking experiments were carried out on ice or in a 4 °C, because the UV lamp generates a substantial amount of heat. Regardless, the irradiation itself also results in a small amount of volume loss when crosslinking in a PCR strip, which should be taken into account. In our experience 25-50 μL volumes are better to mitigate evaporation.
5. There are commercially available 365 nm LED lamps. While these lamps still generate heat, they do result in higher crosslinking efficiencies. In our experience a 365 nm LED lamp heats up the sample to approximately room temperature when the crosslinking is carried out on the benchtop of a 4 ^°^C cold room.
6. Depending on the position of the crosslinker, it might be beneficial to western blot for the subunit containing the Bpa to verify crosslinking.

## Acknowledgments

We thank all members of the Martin lab for discussion and support.

## Author contributions

S.Y.R. and A.M. conceived the study and designed experiments. S.Y.R. cloned constructs, expressed and purified proteins, and performed biochemical measurements and data analyses. S.Y.R. wrote the manuscript with comments from A.M..

## References

1. Bard, J.A.M., Goodall, E.A., Greene, E.R., Jonsson, E., Dong, K.C., Martin, A., 2018. Structure and Function of the 26S Proteasome. Annual Review of Biochemistry 87, 697–724. 10.1146/annurev-biochem-062917-011931

2. Groll, M., Ditzel, L., Löwe, J., Stock, D., Bochtler, M., Bartunik, H.D., Huber, R., 1997. Structure of 20S proteasome from yeast at 2.4Å resolution. Nature 386, 463–471. 10.1038/386463a0

3. Lander, G.C., Estrin, E., Matyskiela, M.E., Bashore, C., Nogales, E., Martin, A., 2012. Complete subunit architecture of the proteasome regulatory particle. Nature 482, 186–191. 10.1038/nature10774

4. Beckwith, R., Estrin, E., Worden, E.J., Martin, A., 2013. Reconstitution of the 26S proteasome reveals functional asymmetries in its AAA+ unfoldase. Nature structural & molecular biology 20, 1164. 10.1038/nsmb.2659

5. Husnjak, K., Elsasser, S., Zhang, N., Chen, X., Randles, L., Shi, Y., Hofmann, K., Walters, K.J., Finley, D., Dikic, I., 2008. Proteasome subunit Rpn13 is a novel ubiquitin receptor. Nature 453, 481–488. 10.1038/nature06926

6. Shi, Yuan, Chen, X., Elsasser, S., Stocks, B.B., Tian, G., Lee, B.-H., Shi, Yanhong, Zhang, N., de Poot, S.A.H., Tuebing, F., Sun, S., Vannoy, J., Tarasov, S.G., Engen, J.R., Finley, D., Walters, K.J., 2016. Rpn1 provides adjacent receptor sites for substrate binding and deubiquitination by the proteasome. Science 351, aad9421. 10.1126/science.aad9421

7. Bard, J.A.M., Martin, A., 2018. Recombinant Expression, Unnatural Amino Acid Incorporation, and Site-Specific Labeling of 26S Proteasomal Subcomplexes, in: Mayor, T., Kleiger, G. (Eds.), The Ubiquitin Proteasome System: Methods and Protocols. Springer, New York, NY, pp. 219–236. 10.1007/978-1-4939-8706-1_15

8. Bard, J.A.M., Bashore, C., Dong, K.C., Martin, A., 2019. The 26S Proteasome Utilizes a Kinetic Gateway to Prioritize Substrate Degradation. Cell 177, 286-298.e15. 10.1016/j.cell.2019.02.031

9. Jonsson, E., Htet, Z.M., Bard, J.A.M., Dong, K.C., Martin, A., 2022. Ubiquitin modulates 26S proteasome conformational dynamics and promotes substrate degradation. Sci Adv 8, eadd9520. 10.1126/sciadv.add9520

10. López-Alfonzo, E.M., Saurabh, A., Zarafshan, S., Pressé, S., Martin, A., 2023. Substrate-interacting pore loops of two ATPase subunits determine the degradation efficiency of the 26S proteasome. 10.1101/2023.12.14.571752

11. Htet, Z.M., Dong, K.C., Martin, A., 2024. The deubiquitinase Rpn11 functions as an allosteric ubiquitin sensor to promote substrate engagement by the 26S proteasome. 10.1101/2024.10.24.620116

12. Chin, J.W., Martin, A.B., King, D.S., Wang, L., Schultz, P.G., 2002a. Addition of a photocrosslinking amino acid to the genetic code of Escherichia coli. Proceedings of the National Academy of Sciences 99, 11020–11024. 10.1073/pnas.172226299

13. Chin, J.W., Santoro, S.W., Martin, A.B., King, D.S., Wang, L., Schultz, P.G., 2002b. Addition of p-Azido-l-phenylalanine to the Genetic Code of Escherichia coli. J. Am. Chem. Soc. 124, 9026–9027. 10.1021/ja027007w

14. Dorman, G., Prestwich, G.D., 1994. Benzophenone Photophores in Biochemistry. Biochemistry 33, 5661–5673. 10.1021/bi00185a001

